# eIF4E3 drives translation of viral mRNAs with short 5’UTRs

**DOI:** 10.64898/2026.05.25.727655

**Authors:** Autumn O. Grimins, Mollie A. Brekker, Sarah Ruth Shick, Ellen L. Suder, Victoria A. Kleiner, Sami Gritli, Ryan Johnston, Alan Wacquiez, Mohsan Saeed, Assen Marintchev, Elke Mühlberger, Rachel Fearns, John Connor, Shawn M. Lyons

## Abstract

Eukaryotic cells express three paralogs of the cap-binding protein eIF4E, yet the functions of the less-studied family members remain poorly understood. Here we demonstrate that eIF4E3, a paralog whose expression is tissue-restricted and whose activity is insensitive to inhibition by eIF4E-binding proteins (4EBPs), drives efficient translation of viral mRNAs with short 5’ untranslated regions (UTRs). Many negative-strand RNA viruses (NSVs), including vesicular stomatitis virus (VSV), respiratory syncytial virus (RSV), and influenza A virus (IAV), produce mRNAs with extremely short 5’UTRs that are incompatible with canonical cap-dependent scanning translation initiation. Using an auxin-inducible degron (AID) system to acutely deplete endogenous eukaryotic initiation factors during active viral infection, we demonstrate that translation of these short-UTR viral mRNAs occurs independently of eIF4E1, the canonical cap-binding protein, while remaining dependent on eIF4E3. In contrast, Ebola virus (EBOV), whose mRNAs bear long, structured 5’UTRs, remains eIF4E1-dependent, implicating 5’UTR length and structural complexity as *cis* determinants of eIF4E paralog selectivity. During VSV infection, 4EBP dephosphorylation sequesters eIF4E1 and broadly suppresses host cap-dependent translation. Because eIF4E3 escapes 4EBP-mediated regulation and preferentially engages short, unstructured 5’UTRs, it is uniquely positioned to sustain viral protein synthesis under these conditions. These findings reveal that VSV exploits a cellular paralog-switching mechanism by co-opting eIF4E3 to maintain viral translation when canonical eIF4F activity is suppressed and establish eIF4E3 as a proviral factor whose tissue-restricted expression in the lung may influence susceptibility to clinically important respiratory pathogens.

---

Translation initiation is the most highly regulated step in protein synthesis, relying on at least 23 individual polypeptides, discounting those within the ribosome, within several individual eukaryotic initiation factors (eIFs) ^1^. Canonical recruitment to a mRNA depends upon the m^7^GTP moiety at the 5’ end of all cellular mRNAs ^2–5^. This 5’ cap is bound by eIF4F^6^, a heterotrimeric complex composed of eIF4E, the cap binding protein^7,8^, eIF4G, a large scaffolding protein^9^ and eIF4A, an ATP-dependent helicase^10^. Interactions between eIF4G and another multisubunit complex called eIF3 serve to recruit the small ribosomal subunit to an mRNA, loaded onto the 40S ribosome as the 43S pre-initiation complex (PIC), which is composed of the 40S ribosome, eIF1/1A, eIF2, eIF3 and eIF5. Recruitment of the 43S PIC to an eIF4F-bound mRNA forms the 48S PIC which then must scan in a 5’ to 3’ direction along the mRNA to find the start codon^11^, which is mediated by several eIFs^12^. Start codon recognition triggers 48S closure and commitment to initiation at a particular site.

The 5’ untranslated region (UTR) of an mRNA serves as a platform for this process. mRNAs from cellular genes have a mean length of 218 nucleotides ^13^. 5’UTR structure and nucleotide composition are major drivers of translation efficiency of an mRNA. Key structural features such as upstream open reading frames (uORFs), secondary structures (e.g., hairpins and G-quadruplexes) or sites for RNA binding proteins have dramatic effects on the initiation rates at the start codon. More generally, the length of a UTR can also affect the translation efficiency of an mRNA. The large 48S PIC necessitates a minimal biophysical length of 15 nucleotides to accommodate mRNA binding ^14^. However, molecular biology experiments have determined that mRNAs with 5’ UTRs of less than 30 nucleotides are poorly translated ^15^. Certain cellular mRNAs, such as histone H4 mRNAs, possess similarly short 5’UTRs ^16^. These mRNAs have adopted a unique mode of cap-assisted translation initiation that promotes efficient translation despite this constraint. Other cellular mRNAs with short 5’UTRs possess special elements known as “translation initiation on short UTRs (TISU)” elements that allow a scanning independent mode of initiation^17^.

The properties of the 5’ UTRs of cellular mRNAs differ from those of the negative strand RNA virus (NSV) mRNAs. The NSVs are major human pathogens, some have non-segmented (nsNSV) genomes, such as rabies virus (RABV), measles virus, and Ebola virus (EBOV), or segmented genomes, such as influenza viruses. Regardless of genome architecture, all NSVs produce capped mRNAs, the majority of which also contain poly(A) tails, thereby fully mimicking cellular mRNAs. However, despite closely mimicking a cellular mRNA, some (but not all) NSV families produce mRNAs that have important distinctions from cellular mRNAs. In particular, members of the families *Rhabdoviridae* (e.g. RABV), *Pneumoviridae* (e.g. respiratory syncytial virus), and *Orthomyxoviridae* (e.g. influenza viruses) generate mRNAs with short 5’ UTRs. For example, the average 5’ UTR length of rhabdovirus mRNAs is only 13 nucleotides. In these cases, cap-dependent assembly of a 48S PIC on this mRNA would position the start codon upstream of the decoding center, precluding start codon recognition^1^. These mRNAs do not possess TISU elements and no stem-loops necessary for a cap-assisted mode of initiation have been described. Despite this constraint, these mRNAs are efficiently translated during infection, raising the question: How do they bypass this restriction?

Some of the most utilized methods in molecular biology are not feasible or optimal for the study of translation. Protein synthesis is amongst the most essential processes in cells^18^. Thus, siRNA-mediated knockdown of factors involved in protein synthesis limits cellular viability more than any other class of factors, including those required for DNA replication. Unsurprisingly, genetic ablation of these factors using CRISPR-mediated approaches is not possible due to their essential nature.

These difficulties are compounded in the study of viral infection. Recapitulating physiological infection kinetics in cells with reduced viability, due to reduction or absence of essential factors, is rarely possible. To circumvent these problems, we have assembled an auxin-induced degron (AID) toolbox of tagged eIFs ^19^. This allows for depletion of endogenously tagged proteins within hours, rather than days, preventing a compensatory cellular response. We have used this approach to reveal principles of mRNA translation of vesicular stomatitis virus (VSV), a member of the rhabdovirus family. VSV has served as the prototypical nsNSV for more than 60 years and has been one of the most intensively studied viruses, leading to key discoveries regarding viral entry, replication, and pathogenesis ^20^. It is also a key reagent in biotechnology, being utilized in oncolytic viral therapies, lentivirus production, and vaccine development. Using this approach, we demonstrate that VSV, and other NSVs with short 5’UTRs, bypass canonical modes of translation initiation by utilizing specific paralogs of eIFs to drive efficient translation of their mRNAs.

## RESULTS

### Generation of a system for rapidly depletion of endogenous eukaryotic initiation factors

To bypass the problem of compensation upon knockdown of a target protein or reduced cell viability that would further confound results, we sought to adapt the auxin inducible degron system from *Arabidopsis thaliana* (**Supp. Figure 1A**). Previous versions of AID technology derived from *Oryza sativa* were unsuitable for studying eIFs due to the high degree of basal activity of the F-box protein, *Os*Tir1; although, recent adaptations have reduced this activity^21^. In contrast, *A. thaliana* AFB2 has been shown to have remarkable low background activity ^19^. Regardless, to reduce the possibility of constitutive activity, in HAP1 cells, we placed *At*AFB2-mCherry under doxycycline (Dox) inducible control so that timing of its expression could be controlled. We titrated doxycycline such that minimal concentrations were needed to fully induce AFB2 expression (**Supp. Figure 1B – C**). We confirmed that neither treatment with dox or auxin (Aux) nor expression of AFB2 perturbed global protein synthesis (**Supp. Figure 1D - E**). Thus, the minimal components necessary for adapting AID-technology for studying cellular protein synthesis are compatible.

Next, we sought to systematically tag eukaryotic initiation factors in Tet-On-AFB2-mCherry HAP1 cells. Using CRISPR/Cas9-mediated homology directed repair (HDR), we integrated AID and epitope (Flag or EGFP) tags into the genomic loci for the *eIF2*β*, eIF4E1, eIF5* and *PABPC1* genes (**Figure 1A - B**). Tagging was confirmed by genotyping PCR and evidenced by mobility shift on SDS-PAGE gels and reactivity with Flag (eIF2β, eIF4E1, eIF5) or GFP (PABPC1) antibodies. Importantly, tagging of individual eIFs did not perturb those proteins found in multisubunit complexes (e.g. eIF4E1 & eIF2β). Further, tagging of these essential proteins did not alter basal protein synthesis indicating that the tags do not interfere with protein function (**Supp. Figure 1F**).

**Figure 1.**
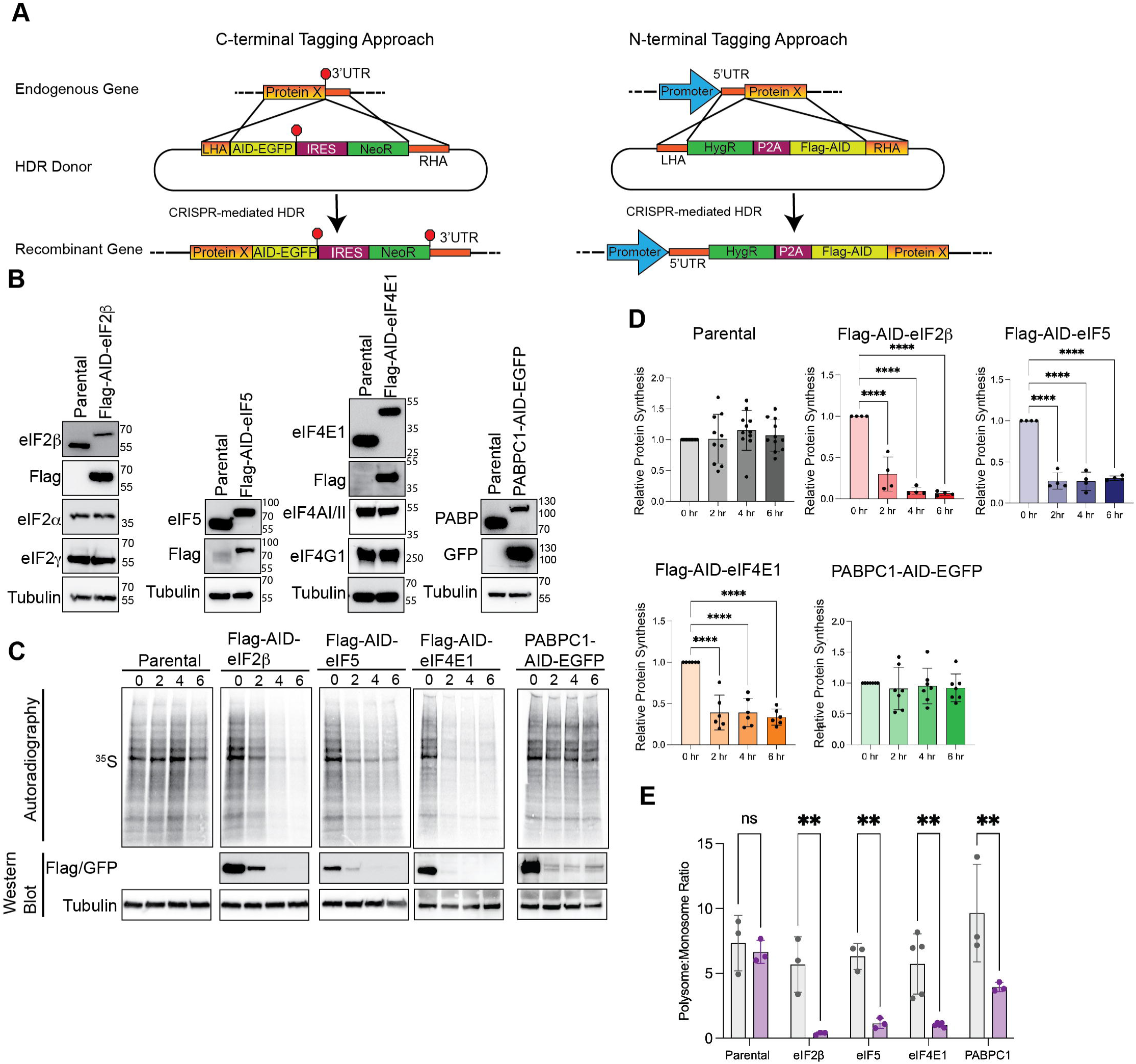
Generation of an auxin-inducible degron (AID) system for eukaryotic initiation factors (eIFs). (**A**) CRISPR/Cas9 mediated homology directed repair (HDR) strategy for insertion of C-terminal (PABPC1) or N-terminal (eIF2β, eIF4E1, eIF5) AID tags. Following drug selection with either G418 (PABPC1) or Hygromycin (eIF2β, eIF4E1, eIF5), single cell clones were isolated. (**B**) Western blot validation of Flag-AID-eIF2β, Flag-AID-eIF4E1, Flag-AID-eIF4E1 and PABPC1-AID-EGFP HAP1 cells. Note mobility shift of tagged protein and reactivity with incorporated epitope (Flag or GFP). Tagging does not perturb other members of those proteins found in multisubunit complexes (eIF2β and eIF4E1). (**C – D**) Addition of auxin triggers rapid depletion of AID tagged protein as monitored by western blotting and inhibition of protein synthesis as monitored by ^35^S-metabolic labeling. Autoradiography was quantified and normalized to total protein from BioRad Stain-free gels (**D**). (**E**) Quantification of polysome-to-monosome (P/M) ratio following 4 hour depletion of indicated protein. Statistical significance determined by one-way ANOVA with Dunnett’s correction for multiple comparisons (**D**) or student’s t-test (**E**). **, p<0.01; ****, p<0.001.

We next sought to determine the depletion kinetics of AID-tagged proteins. Individual cell lines were treated with 100 μg/mL auxin for 2, 4 and 6 hours. Ten minutes prior to harvest, cells were pulsed with [^35^S]-methionine/cysteine to monitor nascent protein synthesis. Lysates were harvested, resolved on SDS-PAGE gels and subjected to autoradiography and western blotting. In each case, AID-tagged protein levels were reduced within 2 hours, with depletion to undetectable levels achieved by 4 hours post-auxin treatment (**Figure 1C**). We confirmed as before that auxin treatment has no effect on protein synthesis in cells lacking an AID-tagged protein (Parental) (**Figure 1C – D**). In contrast, depletion of eIF2β, eIF4E1, and eIF5 sharply reduced cellular protein synthesis within 2 hours post-auxin treatment, albeit to varying degrees with eIF2β depletion having the greatest effect, followed by eIF5 and then eIF4E1. In contrast, acute depletion of PABPC1 had minimal effect on cellular protein synthesis within our time-course, consistent with data from our lab ^22^ and others demonstrating that the primary role of PABPC1 is through mRNA stabilization^23–25^ and that effects of its loss are only evident upon chronic depletion. Via an orthogonal approach, we confirmed inhibition of protein synthesis at 4 hours post-auxin treatment via polysome analysis using sucrose gradient ultracentrifugation (**Figure 1E, Supp. Figure 2A – E**). As with [^35^S]-metabolic labeling, auxin treatment alone had no effect on polysome-to-monosome (P/M) ratios in parental cells, while affecting translation in cells harboring AID-tagged proteins. Thus, we have confirmed the AIDs are suitable for the study of protein synthesis, they achieve rapid depletion of endogenously tagged initiation factors with equally rapid effects on cellular protein synthesis.

### Translation of vesicular stomatitis virus (VSV) mRNAs occurs independently of eIF4E1

VSV is a member of the order Mononegavirales, whose members share a single non-segmented negative-sense genome in 3’ to 5’ orientation. The VSV genome contains only five genes: *nucleoprotein* (N), *phosphoprotein* (P), *matrix protein* (M), *glycoprotein* (G), and *large protein* (L). Each is transcribed as a discrete, monocistronic mRNA bearing a 5’ m^7^G cap and a 3’ poly(A) tails. In this respect, VSV mRNAs closely resemble their cellular counterparts. However, they possess several features that distinguish them from canonical cellular mRNAs. During VSV infection, host protein synthesis is nearly completely inhibited while viral protein synthesis proceeds normally ^26,27^. Other viruses that suppress host translation with comparable efficiency, such as poliovirus^28^, often possess atypical RNA elements, such as internal ribosome entry sites (IRESes)^29,30^, that allow ribosomes to initiate translation in a cap-independent manner. The architecture of VSV mRNAs has long supported the assumption that they are translated by the canonical, cap-dependent mechanism. Yet if host cap-dependent translation is severely restricted during infection, how VSV mRNAs are efficiently translated remains an open question.

Several lines of evidence suggest that VSV mRNAs utilize a specialized route for translation initiation. RPL40, a component of the 60S ribosomal subunit, is dispensable for cellular protein synthesis but required for efficient VSV protein synthesis^31^, hinting at a non-canonical mechanism. Consistent with this, siRNA-mediated knockdown of eIF4G had little effect on VSV translation^32^. Most strikingly, inhibition of host protein synthesis during VSV infection occurs concurrently with dephosphorylation of 4EBP1^33^, which impairs eIF4F complex assembly, suggesting that VSV mRNAs can be translated under conditions where canonical cap-dependent initiation is compromised. The short 5’UTRs of VSV mRNAs reinforce this interpretation: traditional eIF4F-dependent recruitment of the 43S PIC followed by scanning would position the start codon upstream of the decoding center, precluding accurate initiation (**Figure 2A**).

**Figure 2.**
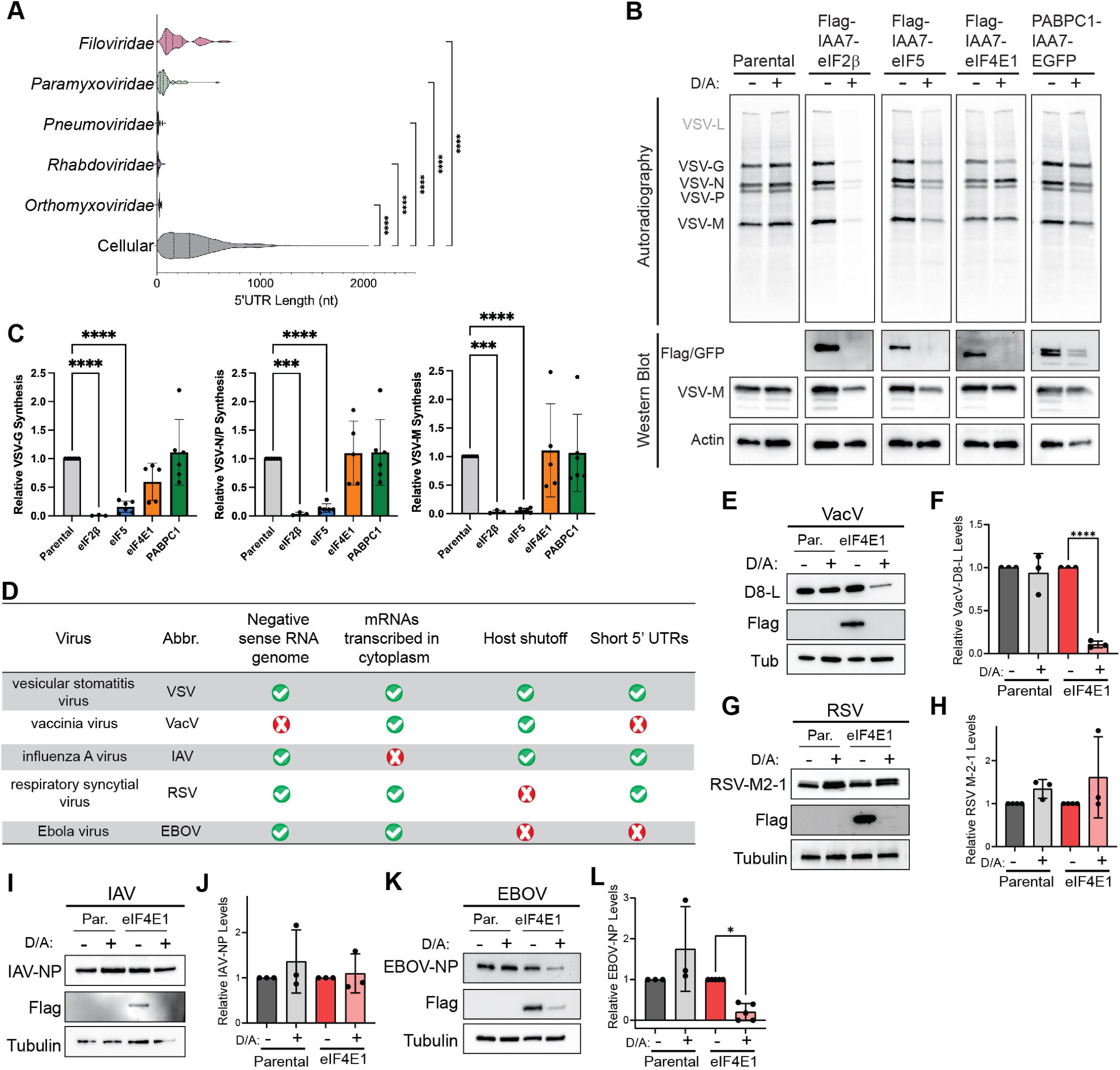
Select negative strand RNA viruses (NSVs) produce mRNAs that are capable of translation in the absence of eIF4E1. (**A**) Analysis of 5’UTR lengths of cellular mRNAs and from select NSV families: *Orthomyxoviridae* [e.g. Influenza A Virus (IAV), Influenza B Virus (IBV)], *rhabodoviridae* [e.g. rabies virus (RABV), vesicular stomatitis virus (VSV)], *Pneumoviridae* [e.g. respiratory syncytial virus (RSV), human metapneumovirus (HMPV)], *Paramyxoviridae* [e.g. measles virus (MeV), Nipah virus (NiV)], *Filoviridae* [e.g. Ebola virus (EBOV), Marburg virus (MARV)]. NSVs mRNAs have significantly shorter 5’UTRs than cellular mRNAs with rhabdovirus, pneumovirus and orthomyxovirus 5’UTRs being especially short. (**B - C**) VSV protein synthesis occurs independently of eIF4E1. Indicated AID-tagged HAP1 cells were depleted of target protein for 2 hours before infection with VSV (MOI=10). Cells were harvested 6 hours post-infection following a brief metabolic labeling with ^35^S-met/cys. Viral proteins were assayed by autoradiography as indicated. Western blotting confirmed depletion of target protein. Viral protein synthesis was quantified based on autoradiography normalized to total protein on biorad stain-free gels (**C**). (**D**) Characteristics of viruses used in this study [VSV, vaccinia virus (VacV), IAV, RSV, EBOV]. (**E - F**) VacV protein synthesis is dependent upon eIF4E1. Cells were depleted of eIF4E1 for 2 hours before infection with VacV. Cells were harvested 24 hours post infection and subjected to western blotting for D8-L, a VacV protein. Loss of eIF4E1 resulted in a failure of D8-L accumulation. (**G – H**) RSV protein synthesis is independent of eIF4E1. Cells were depleted of eIF4E1 for 2 hours before infection with RSV (MOI = 3). Cells were harvested 16 hours post infection and subjected to western blotting for M2-1, a RSV protein. Loss of eIF4E1 did not impair M2-1 accumulation. (**I – J**) IAV protein synthesis is independent of eIF4E1. Cells were depleted of eIF4E1 for 2 hours before infection with IAV (MOI = 3). Cells were harvested 10 hours post infection and subjected to western blotting for IAV NP. Loss of eIF4E1 did not impair NP accumulation. (**K – L**) EBOV protein synthesis is dependent upon eIF4E1. Cells were depleted of eIF4E1 for 2 hours before infection with EBOV (MOI = 3). Cells were harvested 16 hours post infection and subjected to western blotting for EBOV NP. Loss of eIF4E1 resulted in a failure of EBOV-NP accumulation. Statistical significance determined by one-way ANOVA with Dunnett’s correction for multiple comparisons (**C**) or student’s t-test (**F, H, J, L**). *, p<0.05; **, p<0.01; ****p<0.001

To directly test the factor requirements for VSV mRNA translation, we used our AID system to acutely deplete individual initiation factors prior to infection. Cells were depleted for 2 hours, then infected with VSV at MOI = 10. At 6 hours post-infection (8 hours post-auxin treatment), cells were pulsed with [^35^S]-met/cys and viral protein synthesis was assessed by autoradiography (**Figure 2B–C**). As with cellular translation, treatment with dox or auxin alone did not impair viral protein synthesis, and VSV translation remained highly dependent on eIF2 and eIF5 while being unaffected by loss of PABPC1. Strikingly, however, loss of eIF4E1 had no effect on VSV protein synthesis, even as cellular translation was reduced by more than 60%. These data demonstrate that VSV mRNA translation occurs independently of eIF4E1.

We next asked how broadly applicable eIF4E1-independence was among related viruses. To address this, we assembled a panel of viruses with distinct characteristics to systematically define the requirements for eIF4E1-independence (**Figure 2D**). We first validated that our AID system could distinguish eIF4E1-dependent from eIF4E1-independent translation. Vaccinia virus (VacV), a large DNA poxvirus that replicates in the cytoplasm and inhibits host protein synthesis, which are properties shared with VSV, is known to require eIF4F for infection ^32^. AID-mediated depletion of eIF4E1 prevented accumulation of the VacV protein D8-L, confirming that eIF4E1 depletion can block translation of viral mRNAs that had previously been established to require eIF4F (**Figure 2E – F**).

We next tested respiratory syncytial virus (RSV), a nsNSV pneumovirus. Like VSV, RSV produces short, monocistronic mRNAs with short, unstructured 5’UTRs and replicates in the cytoplasm. However, unlike VSV and VacV, RSV does not substantially inhibit host protein synthesis^34,35^. Nevertheless, RSV M2-1 protein continued to accumulate in the absence of eIF4E1 (**Figure 2G – H**), demonstrating that host translational shutoff is neither sufficient, nor necessary for eIF4E1-independence.

We then examined influenza A virus (IAV), a sNSV orthomyxovirus whose mRNAs bear short 5’UTRs, though with some length variability arising from its cap-snatching mechanism of transcription initiation ^36^. IAV inhibits host translation ^37^ but, unlike VSV and RSV, replicates in the nucleus^38^ and its mRNAs must traverse distinct cellular compartments before reaching the cytoplasm for translation. Despite sharing more features with cellular mRNA production and transport, IAV protein accumulation was also unaffected by eIF4E1 depletion (**Figure 2I – J**), consistent with prior RNAi-based evidence^39^ and extending it to an AID system. This result indicates that cytoplasmic replication is not required for eIF4E1-independence.

Finally, we tested EBOV, a filovirus within the nsNSVs. In contrast to VSV, RSV, and IAV, EBOV mRNAs have long, structured 5’UTRs ^40^ more closely resembling those of cellular mRNAs. Accordingly, EBOV protein accumulation was impaired upon eIF4E1 depletion (**Figure 2K – L**). This result suggests that 5’UTR length and structure are key determinants of eIF4E1-independence among negative-strand RNA viruses.

### Short 5’UTRs contribute to eIF4E1-independence

We next sought to investigate whether short 5’ UTRs are a key determinant of eIF4E1-independence. To this end, we took advantage of the well-established reverse genetics system ^41–43^ for VSV to generate recombinant viruses encoding a SNAP-tag reporter inserted between the P and M genes (**Figure 3A**). We rescued two recombinant viruses: one in which the SNAP reporter was preceded by the VSV P mRNA 5’UTR (10 nucleotides, unstructured) and a second in which it was preceded by the EBOV NP 5’UTR (414 nucleotides, highly structured). In both cases, SNAP protein was detectable by metabolic labeling and accumulated with kinetics consistent with other viral proteins, confirming expression from the viral genome (**Supp. Figure 3**). We then infected parental and Flag-AID-eIF4E1 HAP1 cells with each virus in the presence or absence of auxin. SNAP expression driven by the VSV-P 5’UTR retained eIF4E1-independence, mirroring the behavior of authentic VSV mRNAs (**Figure 3B–C**). In contrast, SNAP expression driven by the EBOV NP 5’UTR was reduced, though not completely abolished, upon eIF4E1 depletion. These data agree with our findings that EBOV NP mRNA translation is eIF4E1-dependent and suggest that 5’UTR length and structural complexity act as *cis*-determinants of eIF4E1-independence.

**Figure 3.**
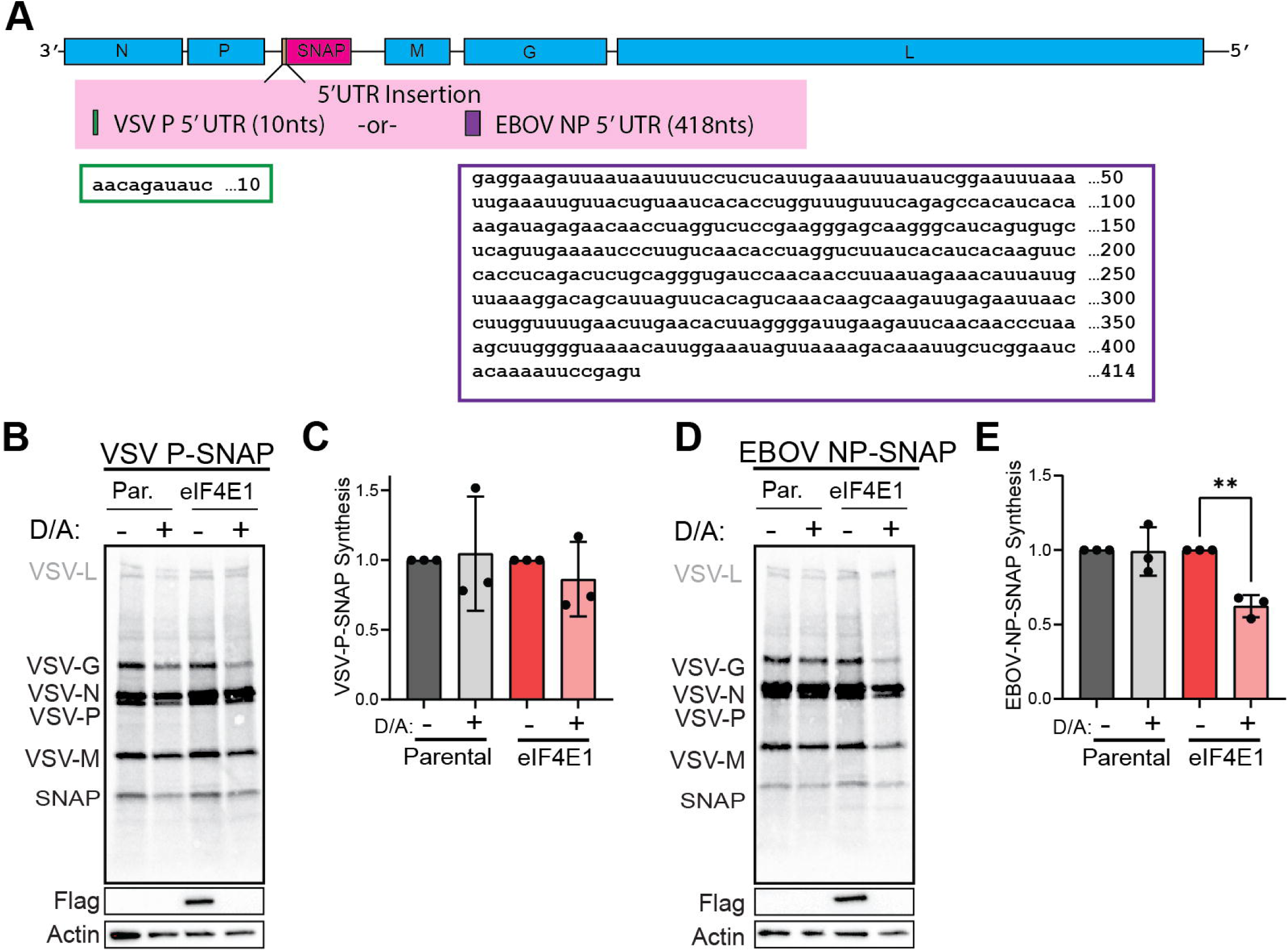
5’UTR length contributes to eIF4E1-independence. **(A)** Schematic of a recombinant VSVs (rVSV) generated. A SNAP gene was inserted between the P and M genes with endogenous VSV gene start and stop signals. The SNAP 5’UTR was derived from either the VSV P mRNA or from the EBOV NP mRNA. Sequence and size of each gene are indicated. (**B - C**) Flag-AID-eIF4E1 cells were infected with rVSV-VSV-P-SNAP (MOI = 10) and harvested 8 hpi following metabolic labeling. Viral proteins and SNAP were analyzed by autoradiography and depletion of eIF4E1 was confirmed by western blotting. As with viral proteins SNAP protein synthesis was unaffected by eIF4E1 depletion. (**D – E**) Flag-AID-eIF4E1 cells were infected with rVSV-EBOV-NP-SNAP (MOI = 10) and harvested 8 hpi following metabolic labeling. Viral proteins and SNAP were analyzed by autoradiography and depletion of eIF4E1 was confirmed by western blotting. SNAP protein synthesis was reduced by ∼50% upon loss of eIF4E1. Statistical significance determined by student’s t-test. **, p<0.01.

### eIF4E3 allow for translation of viral mRNAs with short 5’ UTRs

We next tested the generalizability of these findings by expanding our AID system to HeLa cells. In this background, we incorporated AID tags into the *eIF1*, *eIF4E1*, *eIF4G1*, *eIF5*, and *PABPC1* loci. As in HAP1 cells, editing was confirmed by genotyping and validated by reactivity to epitope tags and by mobility shift of the endogenous proteins (**Supp. Figure 4A - C**). AID-tagged proteins were rapidly depleted upon auxin treatment, with concurrent reductions in protein synthesis observed for eIF4E1 and eIF5 (**Supp. Figure 3D–I**). Depletion of eIF1 did not inhibit protein synthesis. eIF4G1 and PABPC1 cell lines were previously validated^22^, and it was demonstrated that eIF4G1 depletion reduced protein synthesis while PABPC1 depletion had no effect.

Strikingly, in contrast to HAP1 cells, VSV mRNA translation in HeLa cells was dependent upon eIF4E1 (**Figure 4A–B**). As in HAP1 cells, VSV translation was strongly dependent on eIF5 and independent of PABPC1 (**Supp. Figure 5A–D**), and eIF1 was dispensable, consistent with its lack of effect on cellular protein synthesis. Depletion of eIF4G1 also impaired VSV mRNA translation, consistent with a requirement for the intact eIF4F complex in HeLa cells.

**Figure 4.**
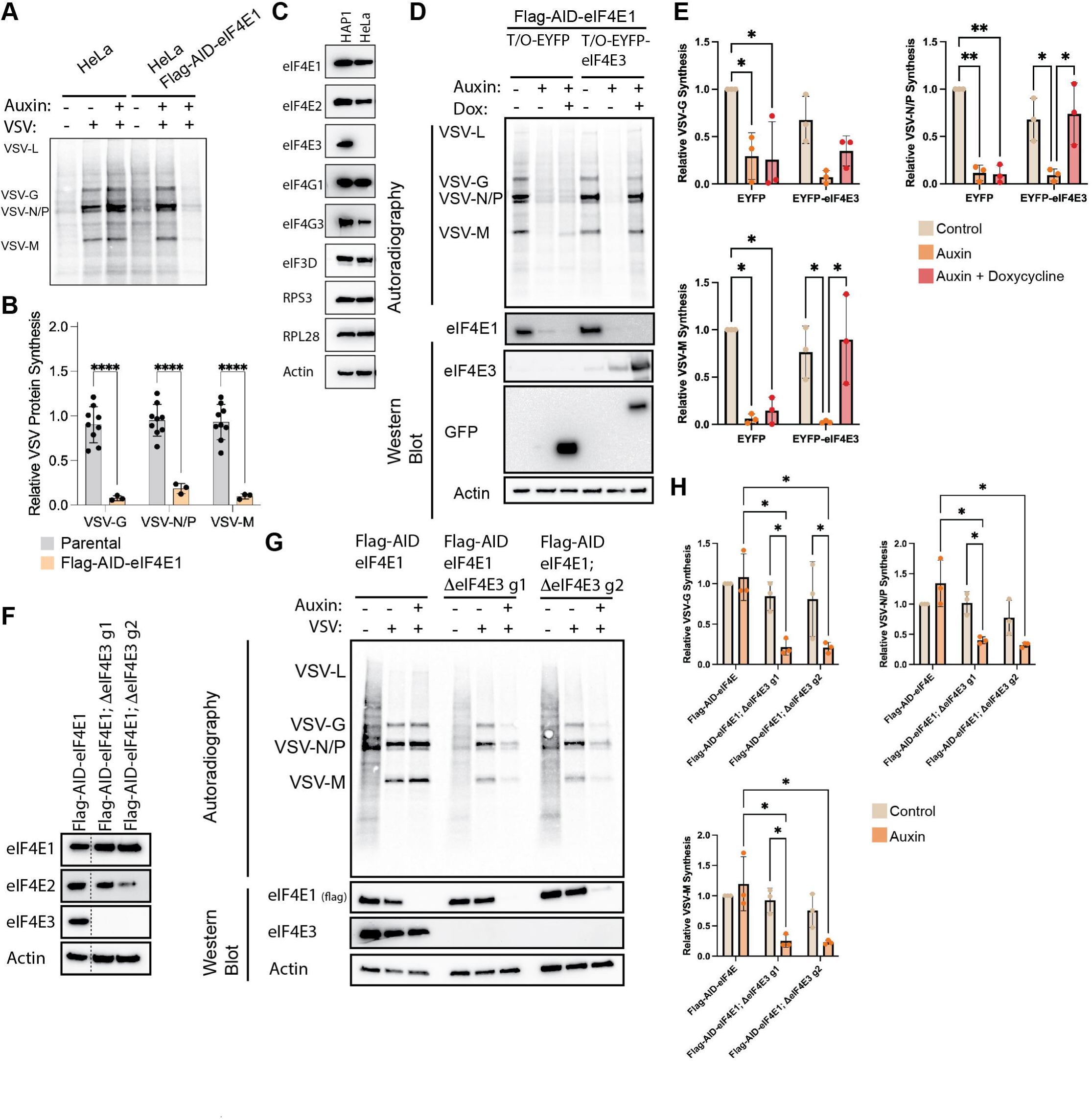
eIF4E3 is necessary and sufficient for eIF4E1-independent translation of VSV mRNAs. (**A - B**) Viral protein synthesis in HeLa cells is dependent upon eIF4E1. Flag-AID-eIF4E1 HeLa cells were depleted of eIF4E1 for 2 hours before infection with VSV (MOI = 10). Cells were harvested at 6 hpi following metabolic labeling with ^35^S-met/cys. In contrast to HAP1 cells, loss of eIF4E1 completely ablated VSV protein synthesis as determined by autoradiography. (**C**) Western blot analysis of HeLa and HAP1 cells reveals that HeLa cells lack expression of eIF4E3, a paralog of eIF4E1, while expression eIF4E2, another eIF4E1 paralog, and eIF3D, a specialized cap-binding subunit of the eIF3 complex. (**D – E**) Exogenous expression of eIF4E3 in HeLa cells rescues eIF4E1-independent expression of VSV proteins. EYFP or EYFP-eIF4E3 was expressed under doxycycline inducible control in Flag-AID-eIF4E1 HeLa cells. Auxin treatment alone triggers loss of endogenous eIF4E1, while co-treatment with auxin and doxycycline triggers loss of endogenous eIF4E1 and exogenous expression of EYFP or EYFP-eIF4E3. Exogenous expression of EYFP-eIF4E3, but not EYFP alone, rescues the eIF4E1-independent expression of VSV proteins as determined by autoradiography. Depletion of eIF4E and expression of EYFP constructs were determined by western blotting. (**F**) Generation of an eIF4E3 knockout Flag-AID-eIF4E1 HAP1 cell line using two individual guide RNAs. (**G – H**) Genetic ablation of eIF4E3 in Flag-AID-eIF4E1 HAP1 cells renders VSV mRNA translation eIF4E1-dependent. Flag-AID-eIF4E1, Flag-AID-eIF4E1;ΔeIF4E3g1 or Flag-AID-eIF4E1;ΔeIF4E3g2 were depleted of eIF4E1 2 hours prior to infection with VSV (MOI = 10) and harvested at 6 hpi. Viral protein synthesis was monitored by autoradiography and depletion of eIF4E1 and eIF4E3 status was validated by western blotting. Statistical significance was determined by student’s t-test (**B**), Two-way ANOVA (**E, H**) *, p<0.05; **,p<0.01; ****, p<0.0001

To understand why VSV mRNA translation differed between HAP1 and HeLa cells, we examined the expression of eIF4E paralogs. Three eIF4E paralogs exist in mammals: eIF4E1, the major cap-binding protein responsible for the bulk of cap-dependent translation; eIF4E2 (also known as 4EHP), which generally functions as a translational repressor^44,45^ though can promote translation in specific contexts^46^; and eIF4E3, a comparatively understudied paralog implicated in translation during stress^47,48^. Comparison of HeLa and HAP1 cells revealed that both express eIF4E1 and eIF4E2, but HeLa cells lack detectable eIF4E3 (**Figure 4C**). Broader analysis of commonly used cell lines revealed variable eIF4E3 expression (**Supp. Figure 6**); notably, while HeLa cells express no detectable eIF4E3, HEp-2 cells, a subclone of HeLa, do, suggesting that eIF4E3 expression is context-dependent. This agrees with previous work that has shown that eIF4E3 mRNA expression is tissue restricted, with highest levels in the heart, lung and skeletal muscle^49^. These observations led us to hypothesize that eIF4E3 is required for eIF4E1-independent VSV mRNA translation in HAP1 cells. To test this, we generated HeLa Flag-AID-eIF4E1 cells stably expressing doxycycline-inducible EYFP or EYFP-eIF4E3. In this system, auxin treatment depletes eIF4E1, while co-treatment with auxin and doxycycline depletes eIF4E1 and simultaneously induces eIF4E3 expression (**Figure 4D**).

Consistent with our earlier findings, depletion of eIF4E1 in HeLa cells strongly reduced VSV protein synthesis, and this defect was not rescued by expression of EYFP alone. Strikingly, expression of EYFP-eIF4E3 fully restored VSV mRNA translation in the absence of eIF4E1, demonstrating that eIF4E3 is sufficient to support VSV translation when eIF4E1 is unavailable (**Figure 4D–E**). These data establish eIF4E3 as a bona fide alternative cap-binding protein capable of driving VSV mRNA translation.

To determine whether eIF4E3 is also necessary for eIF4E1-independent VSV translation, we turned to a loss-of-function approach in HAP1 cells, which endogenously express eIF4E3. We generated eIF4E3 knockout HAP1 cells using two independent guide RNAs to control for potential off-target effects of CRISPR/Cas9 editing (**Figure 4F**). In cells lacking eIF4E3, depletion of eIF4E1 led to a significant reduction in VSV protein synthesis, phenocopying the dependence on eIF4E1 observed in HeLa cells (**Figure 4G–H**). Together, these gain- and loss-of-function data demonstrate that eIF4E3 is both sufficient and necessary for eIF4E1-independent translation of VSV mRNAs, establishing it as a functional paralog that supports viral protein synthesis when canonical cap-dependent initiation is compromised. Notably, eIF4E3 has been shown to be refractory to inhibition by 4EBPs^47,49,50^, which sequester eIF4E1 during stress and viral infection, and has been implicated in preferential translation of mRNAs with short 5’UTRs^47^. These properties may explain its specific role in supporting VSV mRNA translation under conditions where canonical eIF4F activity is compromised.

## DISCUSSION

The study of essential cellular processes presents a fundamental methodological challenge: the factors most important to understand are often those whose disruption is most toxic to cells. This is acutely true for translation initiation, where genetic ablation of eukaryotic initiation factors is not feasible and siRNA-mediated depletion is confounded by incomplete knockdown and severe reductions in cellular viability. Here, we demonstrate that auxin-inducible degron technology, adapted from *A. thaliana* AFB2 ^19^, overcomes these limitations for the study of translation. AID-mediated depletion achieves complete loss of target protein within hours, a timescale that elicits clear translational consequences while minimizing secondary effects. This provides a critical advantage over siRNA-based approaches that require days to achieve comparable depletion. By coupling AID with a doxycycline-inducible expression system, we have established a rapid “deplete-and-replace” platform that we have leveraged previously to dissect translational regulation and extend here to the study of host-virus interactions.

Other degron-based approaches have been applied to the study of mRNA metabolism in human cells. Notably, the Fabian lab achieved PABP depletion using a DHFR-based degron system ^24^, which, while effective, operates on substantially longer decay kinetics than the AID system described here. The shorter timescale of AID-mediated depletion is particularly valuable in the context of translation regulation, where compensatory responses to factor loss can confound interpretation within hours. Importantly, this platform is not limited to the experimental design employed here. The ability to deplete a target protein at a defined point during viral infection opens new experimental possibilities. For example, poxviruses require eIF4E for early gene expression^32^, but published data suggest that late gene expression may be eIF4E-independent^51^. The AID system described here would allow eIF4E to be removed at defined intervals post-infection, enabling direct temporal dissection of factor requirements across the viral lifecycle, which had not previously been possible.

An important caveat to AID-based approaches is that endogenous tagging of essential proteins requires careful validation. If an epitope or degron tag perturbs protein function, the resulting cell line will itself have translational defects independent of auxin treatment, which will confound all subsequent experiments. We addressed this by confirming that AID-tagged proteins retain normal complex assembly and that tagged cell lines exhibit wild-type levels of basal protein synthesis prior to auxin addition. Nevertheless, not all tagging attempts were successful. For example, our initial efforts to tag eIF4E1 at its C-terminus failed to yield fully edited cell lines. This is consistent with the C-terminus of eIF4E1 being adjacent to the m^7^GTP binding domain, suggesting that C-terminal tags may sterically interfere with eIF4F complex activity. In contrast, the N-terminal region does not play a role in eIF4G interaction or m^7^GTP binding. We also attempted to generate AID-tagged cell lines for additional factors, including eIF2α and eIF1A, without success. Whether these failures reflect the functional importance of the targeted termini or simply inefficient guide RNA activity at those loci remains unclear. Collectively, across two cell lines we successfully generated six AID-tagged proteins, eIF1, eIF2β, eIF4E1, eIF4G1, eIF5, and PABPC1, each of which passed functional validation prior to use.

This study also highlights a key role for eIF4E paralogs in cellular biology. As the major cap-binding protein, eIF4E1 has been intensively studied since its discovery, and its regulation by 4EBPs and MNK kinases is well characterized. eIF4E2/4EHP has similarly received considerable attention, particularly in the context of miRNA-mediated translational repression via the 4EHP-GIGYF2 complex ^52^, though it also promotes translation under hypoxic conditions through assembly of an alternative eIF4F complex with eIF4G3^46,53^. By contrast, remarkably little is known about eIF4E3. This likely reflects its restricted and highly variable expression across cell types, a point underscored by our own findings that eIF4E3 is readily detectable in HAP1 but absent in HeLa cells. Most prior studies of eIF4E3 have relied on transient overexpression or biophysical approaches, yielding conflicting conclusions about whether it functions as a translational activator or repressor. The Borden lab demonstrated that eIF4E3 binds the m^7^G cap via an atypical single-tryptophan mechanism with 10 - 40-fold lower affinity than eIF4E1, and that overexpressed eIF4E3 suppresses translation of eIF4E1 target mRNAs, suggesting a tumor suppressive function through competitive cap binding ^50^. In contrast, Landon et al. demonstrated that eIF4E3 can form a functional eIF4F-like complex with eIF4G and eIF4A and drives translation of a distinct subset of mRNAs ^48^. These observations were reconciled by Weiss et al., who demonstrated that under basal conditions endogenous eIF4E3 does not assemble into an eIF4F complex because eIF4E1 outcompetes it for the available eIF4G pool^47^. However, when eIF4E1 is sequestered by hypophosphorylated 4EBPs, as occurs during VSV infection^33^, eIF4E3 assembles into an alternative eIF4F complex and preferentially drives translation of mRNAs with short, unstructured 5’UTRs. Our findings extend this model to a new biological context: viral infection, where VSV and other negative-strand RNA viruses with short 5’UTRs exploit precisely this eIF4E3-dependent mechanism to sustain viral protein synthesis under conditions that broadly suppress eIF4E1-mediated translation. In doing so, this work establishes eIF4E3 as a physiologically important translational regulator whose activity is revealed not in unstressed cells but in a context where canonical cap-dependent translation is compromised.

During VSV infection, 4EBP dephosphorylation occurs alongside the action of VSV matrix protein as a dual mechanism driving host translational shutoff ^33,54,55^. As 4EBPs accumulate in their active, dephosphorylated state, eIF4E1 is progressively sequestered, broadly suppressing cap-dependent translation of cellular mRNAs. Because eIF4E3 escapes this regulation and is preferentially attuned to the short, unstructured 5’UTRs characteristic of VSV mRNAs, it is well positioned to selectively sustain viral protein synthesis under these conditions. Together, these observations suggest a model in which VSV utilizes both eIF4E1 and eIF4E3 early in infection, before 4EBP dephosphorylation is fully established, but becomes increasingly reliant on eIF4E3 as host translational suppression deepens over the course of infection. Directly testing this temporal model would require kinetic analysis of eIF4E paralog usage at distinct stages of infection, which remains an important avenue for future investigation.

VSV mRNAs, by virtue of their short unstructured 5’UTRs, are precisely the class of transcripts that eIF4E3 preferentially engages, making them ideal substrates for eIF4E3-dependent translation under conditions of eIF4E1 suppression. Unlike eIF4E1 and eIF4E2, whose expression is ubiquitous, eIF4E3 expression is tissue-restricted^49^, with particularly high levels reported in lung tissue. This is notable given that RSV and IAV, two of the short-UTR NSVs we demonstrate are eIF4E1-independent, are primary respiratory pathogens whose principal site of infection is the lung epithelium, suggesting that eIF4E3 expression may be physiologically relevant to infection by these viruses *in vivo*. More broadly, the mechanistic framework established here generates testable predictions for other NSVs not examined in this study. Human metapneumovirus (HMPV), a pneumovirus closely related to RSV, produces mRNAs with similarly short 5’UTRs and also infects the same respiratory epithelium, and would therefore be predicted to exhibit eIF4E1-independence. Measles virus and human parainfluenza viruses (HPIV), paramyxoviruses whose mRNAs bear 5’UTRs that are significantly shorter than those of cellular mRNAs but longer than those of rhabdoviruses, orthomyxoviruses, and pneumoviruses, represent strong candidates for future analysis to define the boundaries at which 5’UTR length restricts eIF4E3 usage. By contrast, filoviruses such as EBOV and Marburg virus (MARV), which bear the long-structured 5’UTRs, possess eIF4E1-dependence (EBOV) or would be predicted to rely on eIF4E1 (MARV) for efficient translation. Intriguingly, this dependence may reflect a broader biological constraint: unlike the respiratory viruses discussed above, which replicate primarily in a single tissue, filoviruses cause systemic infections and must replicate productively across a diverse array of cell types and tissues, many of which may express little or no eIF4E3. Reliance on the ubiquitously expressed eIF4E1, rather than the tissue-restricted eIF4E3, may therefore be an adaptation that enables the broad cellular tropism characteristic of filovirus infection. The 5’UTR length distributions we present in Figure 2A provide a roadmap for such predictions across the broader Mononegavirales order, and directly testing these predictions will be an important direction for future work.

The partial dependence of EBOV NP-SNAP translation on eIF4E1 in our recombinant VSV system warrants consideration. eIF4E3 preferentially promotes translation of mRNAs with short, unstructured 5’UTRs, consistent with its role in supporting VSV mRNA translation described here. However, eIF4E3 retains partial capacity to support translation of mRNAs with longer, more structured 5’UTRs, albeit with reduced efficiency^47^. In our recombinant system, where only the SNAP reporter bears the long, structured EBOV NP 5’UTR, this partial eIF4E3 activity is sufficient to sustain a detectable level of SNAP protein synthesis in the absence of eIF4E1. This interpretation is consistent with a model in which eIF4E3 exhibits a strong but not absolute preference for short UTRs, with 5’UTR length and structural complexity acting as a rheostat governing eIF4E3-dependent translation efficiency rather than an “all-or-nothing” switch. Importantly, this distinction has significant implications for authentic EBOV infection, in which all seven viral mRNAs bear long, structured 5’UTRs. Under these conditions, the reduced efficiency of eIF4E3 on each individual mRNA would be compounded across the entire viral proteome, including the polymerase complex subunits L and VP35 that are essential for viral transcription. The resulting deficit in viral mRNA production would further amplify the translational defect, explaining the stronger eIF4E1-dependence observed during authentic EBOV infection relative to our recombinant system. Together, these data support a model in which eIF4E3 is functionally attuned to short, unstructured 5’UTRs, and in which the uniform presence of long UTRs across all EBOV mRNAs creates a compounded vulnerability to eIF4E1 loss that cannot be fully compensated by eIF4E3 activity alone.

Why eIF4E3 is specifically attuned to translation of mRNAs with short 5’UTRs remains an open and important question, though existing biophysical data offer a starting point for speculation. The atypical mode of cap recognition by eIF4E3 that relies on a single tryptophan rather than the canonical aromatic sandwich likely results in a 48S PIC with a distinct architecture compared to one assembled on an eIF4E1-bound mRNA ^48,50^. This structural difference could alter the effective distance between the cap-proximal face of the PIC and the decoding center, reducing the minimum 5’UTR length required for accurate start codon positioning by shrinking the biophysical “blind spot” that makes very short UTRs poorly translated under canonical conditions. Alternatively, the lower cap affinity of eIF4E3 relative to eIF4E1 may facilitate more rapid dissociation of eIF4E3 from the cap upon 43S PIC loading, potentially enabling the ribosome to be positioned closer to the 5’ end and recognize proximal start codons that would otherwise be occluded. A major unresolved question raised by either model is how selectivity for short UTRs is achieved at the level of initial cap recognition. Since eIF4E3 presumably cannot interrogate UTR length prior to cap binding, selectivity likely emerges post-binding. This could be achieved by differential stabilization of the eIF4F complex on short versus long UTRs, or through preferential retention of eIF4E3-containing complexes at mRNAs whose architecture facilitates rapid scanning completion. Ribosome profiling of eIF4E3-dependent versus eIF4E1-dependent translatomes under matched conditions, combined with structural analysis of eIF4E3-containing 48S complexes, will be necessary to distinguish among these possibilities.

More broadly, these findings suggest that the expression pattern of eIF4E3 across cell types and tissues may be an underappreciated determinant of susceptibility to infection by negative-strand RNA viruses. Cells lacking eIF4E3 would be predicted to rely exclusively on eIF4E1 for translation of short-UTR viral mRNAs and therefore to be more vulnerable to the translational consequences of 4EBP-mediated eIF4E1 suppression during infection. Whether eIF4E3 expression influences viral replication efficiency, cell-type tropism, or disease severity in vivo represents an important and unexplored question.

## MATERIALS AND METHODS

### Cell culture

HAP1, HeLa, Vero E6, MDCK, 293T, and BHK-21 cells were cultured in 1X Dulbecco’s Modified Eagle’s Medium (DMEM) containing 10% fetal bovine serum (Atlas Biologics) and 1X penicillin-streptomycin (Gibco). HEp2 cells were cultured in 1X Opti-MEM Reduced Serum Media containing 2% FBS and 1X pen-strep. All cells were incubated at 37℃ with 5% CO_2_ and routinely screened to ensure no mycoplasma contamination.

### Generation of AID-tagged cell lines

Custom plasmids were generated for N-terminal or C-terminal Homology directed repair constructs (pHDR) containing the IAA7 AID tag^19^ followed by the EGFP ORF, EMCV IRES and a Neomycin resistance cassette (PABPC1) or a hygromycin resistance cassette followed by a P2A self-cleaving peptide, 1x FLAG tag and the IAA7 AID tag (eIF4E1, eIF5, eIF2β) flanked by two multiple cloning sites. Left and right homology arms (LHA and RHA) were cloned into either multiple cloning site comprising approximately 250 bp upstream and downstream of the site of genomic insertion. Plasmid construction was confirmed by whole plasmid sequencing (Plasmidsaurus). All PCR was performed using Phusion or Q5 DNA polymerases (NEB) according to manufacturer’s instructions. All oligonucleotides were synthesized by integrated DNA technologies (IDT). Guide RNAs (gRNAs) were designed using ATUM gRNA design tool using DNA surrounding the start or stop codon (https://www.atum.bio/eCommerce/cas9/input) and cloned into pCas-Guide (Origene). EYFP and EYFP-eIF4E3 were cloned into pPB-TetOne-NeoR. AFB2-mCherry HeLa cells were generated by co-transfecting pSH-EFIRES-P-AtAFB2-mCherry-weak NLS (A gift from Elina Ikonen- Addgene plasmid #129717) with px300-AAVS1 to incorporate the expression cassette into the AAVS1 safe-harbor locus. For HAP1 cells, the AtAFB2-mCherry-Weak NLS construct was subcloned into pLVX-tetone lentiviral vector. Lentiviral particles were generated using 293T cells and HAP1 cells were transduced. Both HAP1 and HeLa cells were selected with puromycin and single cell cloned by limiting dilution according to previously established procedures ^56^.

For incorporation of AID-tags, pCas-guide and pHDR plasmids were co-transfected (1.25 μg each) into HeLa or HAP1 cells grown in antibiotic free media plated in a 6-well tissue culture dish using Lipofectamine 2000 according to manufacturers instructions. Cells were incubated for 24 hours. For HAP1 cells, p53 was inhibited by treatment with pitithrin-α. Cells were selected with appropriate selection reagent (G418 or Hygromycin). Cells were cloned by limiting dilution. Single cell clones were screened by genotyping using Platinum direct PCR mix (Invitrogen) according to manufacturer’s instructions followed by western blotting.

Complete list of plasmids and primers can be found supplemental data.

### CRISPR-Cas9 mediated knockout

gRNAs targeting eIF4E3 were designed using ATUM gRNA design tool using DNA surrounding the start or stop codon (https://www.atum.bio/eCommerce/cas9/input) and cloned into lentiCRISPRv2 Blast (A gift from Brett Stringer-Addgene plasmid #98293). Lentiviral particles were generated using 293T cells and used to transduce HAP1 cells. Cells were selected with blasticidin and single cell clone.

### AID-mediated depletion

HAP1 Tet-On-AFB2-mCh cells were induced with 100 ng/mL of doxycycline (Dox) containing media for 16 hours prior to experimentation. To deplete AID-tagged protein, cells were treated with 100 μg/ml indole-3-acetic acid (auxin, Aux) for indicated times.

### ^35^S-metabolic labeling and western blotting

All labeling experiments were performed using EasyTag EXPRESS ^35^S Protein Labeling Mix, containing radioactive methionine and cysteine. Cells were plated between 70-80% confluence for the day of the experiment. Prior to harvesting, cells were incubated in 1X DMEM, Methionine Free with 10% FBS for 15 minutes followed by incubation with 11μCi/ml ^35^S protein labeling mix in the same media. Cells are then washed twice with 1X PBS prior to protein extraction with NP-40 Lysis buffer (10 mM Tris [pH 8.0], 150 mM NaCl, 0.5% NP-40). Lysates were quantified using bradford colorimetric assay. Equal amounts of lysates were resolved by SDS-PAGE electrophoresis using BioRad stain-free gels. Total protein was imaged and quantified using BioRad ImageLab. Protein was transferred to nitrocellulose membranes, dried, and exposed to phosphor storage screens. Phosphor imaging was performed with the Amersham Typhoon Imager. Any membranes used for further western blot probing were rehydrated in 1X TBST prior to blocking in 5% non-fat dry milk/TBST. Primary antibody incubation was performed in 1X TBS supplemented with 5% normal horse serum and 0.2% sodium azide. Secondary antibody incubation was performed in 1X TBS supplemented with 5% normal horse serum. Antibodies were detected by chemiluminescence. Autoradiography and western blotting were quantified using BioRad Image lab and normalized to total protein signal.

### Viral infections

For experiments in combination with AID-mediated depletions, Auxin was added 2 hours prior to infection to ensure infection occurs prior to target protein depletion. All viral stocks were titered prior to experimentation and thawed on ice. VSV and VacC infection inoculum was prepared in 1X DMEM. Cells were infected and at harvested at indicated timepoints. IAV infection inoculum was prepared iPBS as describe in ^57^ (1x DPBS (Mg/Ca^2+^) supplemented with 1x pen-strep, 0.1% v/v FBS, 0.3% BSA). Virus was adsorbed for 1 hour before iPBS was replaced with post-infection DMEM (piDMEM, Dulbecco’s DMEM supplemented with 1x Pen-strep, 0.3% BSA, 0.1% FBS, 20 mM HEPES [pH 7.4], 1 μg/mL TPCK-treated Trypsin). RSV-A2 infection was performed in 1X OPTI-MEM. All EBOV experiments were performed under biosafety-level 4 (BSL4) conditions at the National Emerging Infectious Diseases Laboratories BSL4 suite in Boston, Massachusetts using Zaire EBOV.

### Generation of recombinant vesicular stomatitis virus

Recombinant VSV was prepared based on previously described protocols ^41^. Full length VSV cDNA clone and codon optimized T7 plasmids were generously provided by Drs. Rahm Gummuluru and Adam Hume. A SNAP reporter was cloned in between VSV-P and VSV-M, generating pVSV.2^PSM^ genome plasmid (P-SNAP-M). 5’UTR region of VSV-P and EBOV-NP were amplified by PCR and inserted into pVSV.2^PSM^ to flank the SNAP reporter. Final plasmids were validated by Plasmidsaurus sequencing.

BHK cells were seeded into 6-well tissue culture plates for 80% confluency on the day of transfection. Genome and optimized helper plasmids (T7, VSV-N, VSV-P, and VSV-L) were transfected using Lipofectamine 2000 and incubated for 24 hours. Cells were monitored for cytopathic effect. P1 stocks were collected following the appearance of cytopathic effect. P1 stocks were expanded and titered following the standard protocols. Virus from P2 aliquots was assessed by RT-PCR and amplicon sequencing.

## Supplementary Figure

**Supplementary Figure 1**. **Validation and optimization of HAP1 AID system**. (**A**) Schematic of Tet/On-AFB2-mCherry HAP1 cell function in combination with AID-tagged proteins. Doxycycline induces expression of AFB2-mCherry, while subsequent addition of auxin depletes tagged protein. (**B – C**) Optimization of doxycycline concentrations needed to induce AFB2-mCherry expression. (**D – E**) Neither addition of auxin, nor doxycycline, affects basal protein synthesis as determine by ^35^S-metabolic labeling. Nascent protein synthesis was determined by autoradiography and normalized to total protein from biorad stain-free gel. AFB2-mCherry expression was determine by western blotting. (**F**) Endogenous tagging of indicated eIFs with AIDs does not impair basal protein synthesis rates as determined by ^35^S metabolic labeling.

**Supplementary Figure 2**. **Polysome analysis of indicated AID tagged cells following target protein depletion**. Parental HAP1 (**A**), Flag-AID-eIF2β (**B**), Flag-AID-eIF5 (**C**), Flag-AID-eIF4E1 (**D**), and PABPC1-AID-EGFP (**E**) cells were depleted of AID-tagged proteins for 4 hours. Lysates were prepared and subjected ultracentrifugation (238,000 *x g*, 2 hr) through 10 – 50% sucrose gradients. Gradients were continuously collected and A260 was monitored indicating 40S, 60S, 80S and polysomal peaks. Area under the curve (AUC) was calculated for monosome and polysomes.

**Supplemental Figure 3. Time course of recombinant VSV infection.** Recombinant VSV-WT (**A**), VSV-VSV-P-SNAP (**B**), and VSV-EBOV-NP-SNAP (**C**) viruses were rescued using plasmid-based recovery system in BHK cells. HAP1 cells were infected with individual viruses (MOI = 10). Recombinant VSV containing SNAP genes express SNAP protein as indicated by autoradiograph and by western blotting to SNAP protein. SNAP-containing viruses have slightly delayed kinetics regarding inhibition of host-protein synthesis as is typical upon increasing the length of the viral genome (**D**).

**Supplemental Figure 4**. **Generation of HeLa AID-tagged cell lines**. eIF1 (**A**), eIF5 (**B**), and eIF4E1 (**C**) were tagged with AID and Flag in HeLa cells constitutively expressing AFB2-mCherry. Note the mobility shift of tagged proteins and reactivity with Flag antibody. eIF4G1-AID-Flag and PABPC1-AID-EGFP cells were characterized previously (Johnston et al. 2026). Timecourse following auxin treatment in eIF1-AID-Flag (**D – E**), Flag-AID-eIF5 (**F – G**), and Flag-AID-eIF4E1 (**H – I**) HeLa cells. eIF1 depletion did not negatively affect cellular protein synthesis, although we note a slight, but insignificant increase in protein synthesis, consistent with eIF1’s role in preventing translation until ideal start codon contexts are revealed. Both eIF5 and eIF4E1 depletion resulted in rapid and significant inhibition of protein synthesis as determined by autoradiography and depletion of AID-tagged protein was monitored by western blotting against endogenous protein and Flag epitope. Statistical significance determined by one-way ANOVA with Dunnett’s correction for multiple comparisons. **, p<0.01; ****,p<0.0001.

**Supplemental Figure 5. Requirements for addition eIFs for VSV protein synthesis in HeLa cells.** (**A - B**) VSV protein synthesis depends upon eIF5 in HeLa cells, consistent with findings in HAP1 cells. VSV partially requires eIF4G in HeLa cells. Discordance with eIF4E1 expression is likely due to eIF4G3-dependent buffering (see Johnston et al. 2026). eIF1 depletion slightly increases VSV protein synthesis, consistent with our findings for cellular protein synthesis. Statistical significance determined by one-way ANOVA with Dunnett’s correction for multiple comparisons. *, p<0.05; ****, p<0.0001.

**Supplemental Figure 6. Cell type variability expression of eIF4E3.** Select commonly used human tissue culture cells were assayed for eIF4E3 expression by western blotting. eIF4E3 ranges from not expressed (HeLa) to weakly expressed (BEAS-2B, A549, Hep2, MCF7) to high levels of expression (293T, Calu3, HAP1). Of note is the difference between HeLa and Hep2 cells. Hep2 cells are a derivative of HeLa cells.

