## Supplementary Data for "eIF4E3 drives translation of viral mRNAs with short 5’UTRs"

### Supplementary Figure 1

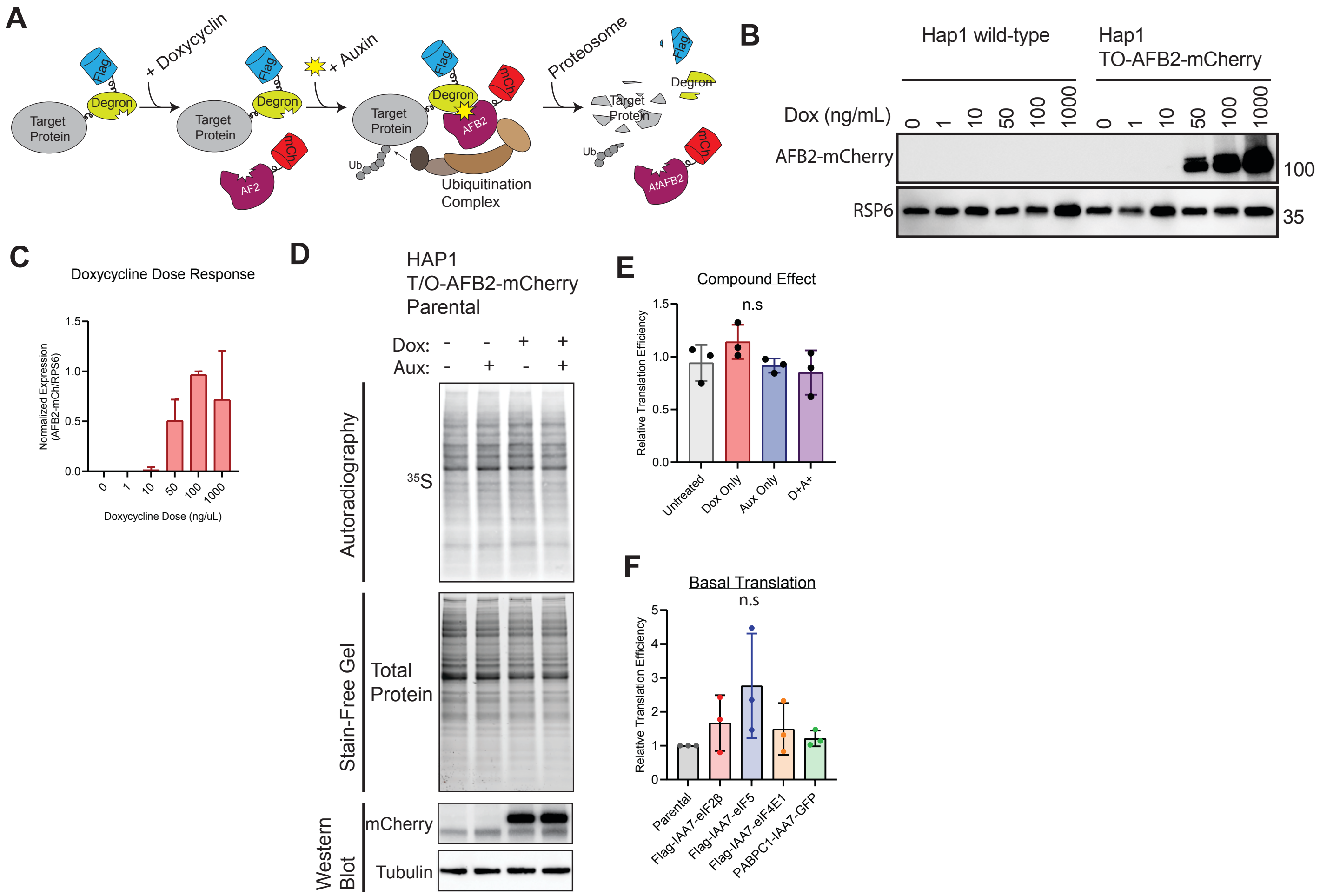

### Supplemental Figure 2

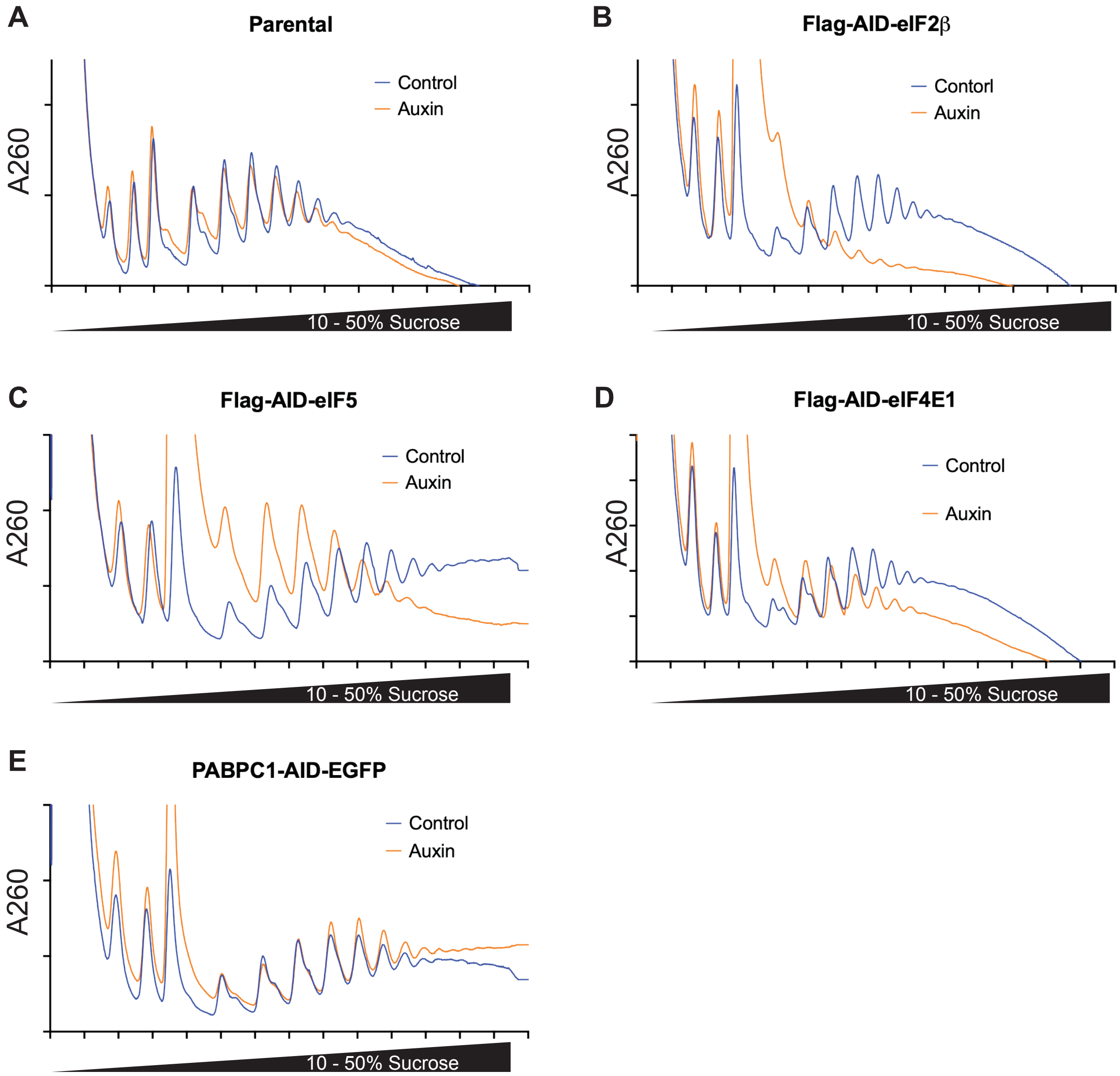

**Supplemental Figure 3**

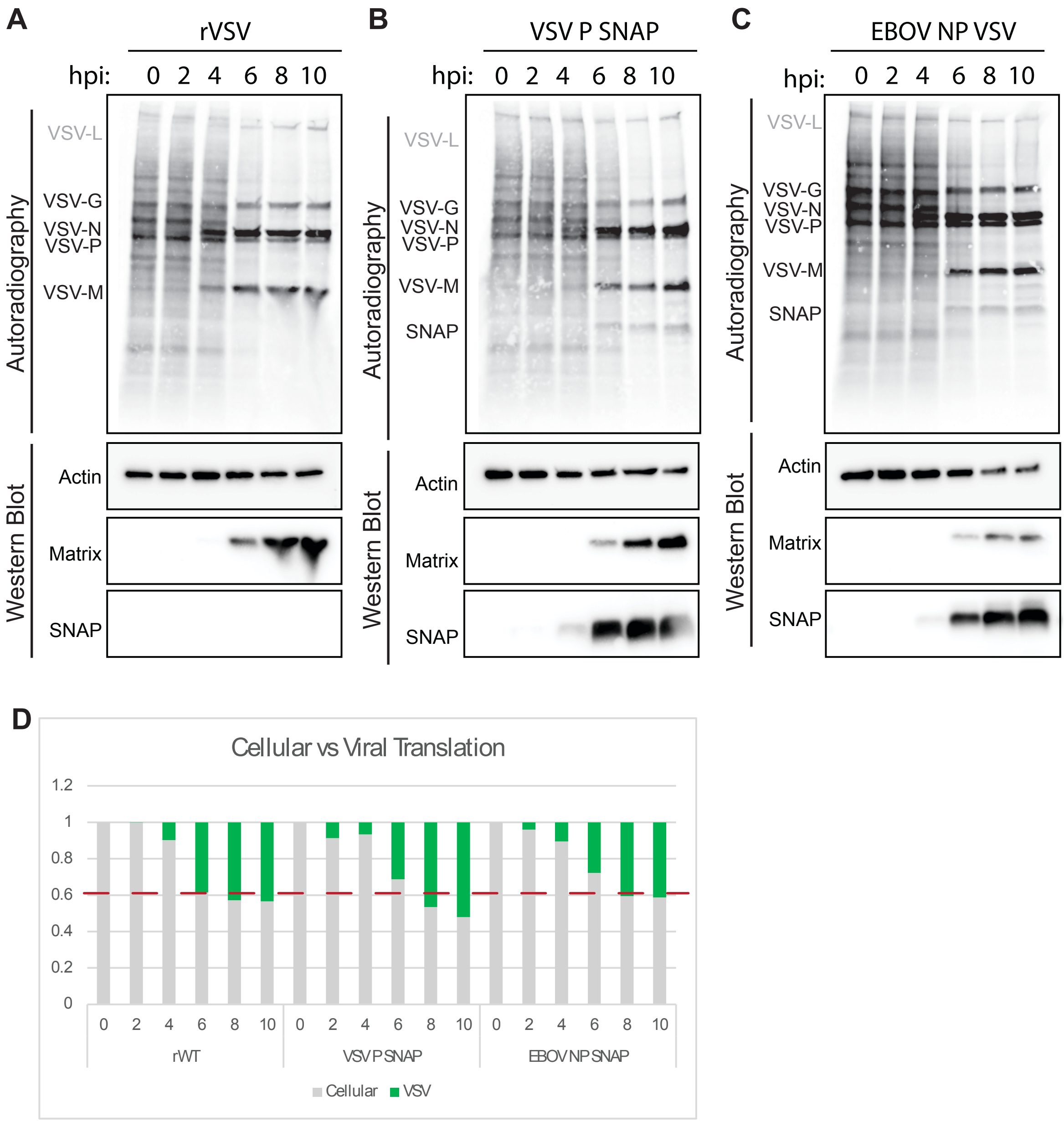

#### Supplemental Figure 4

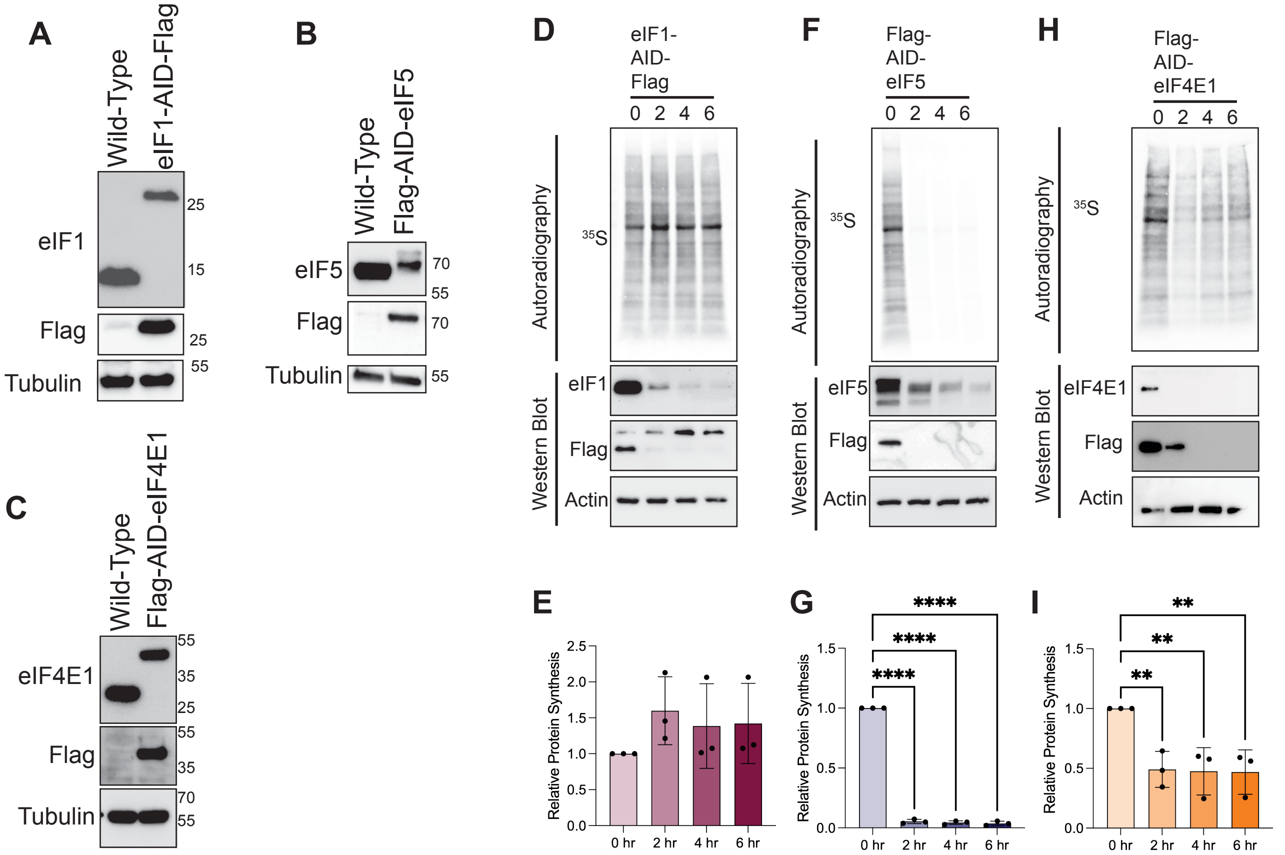

#### Supplemental Figure 5

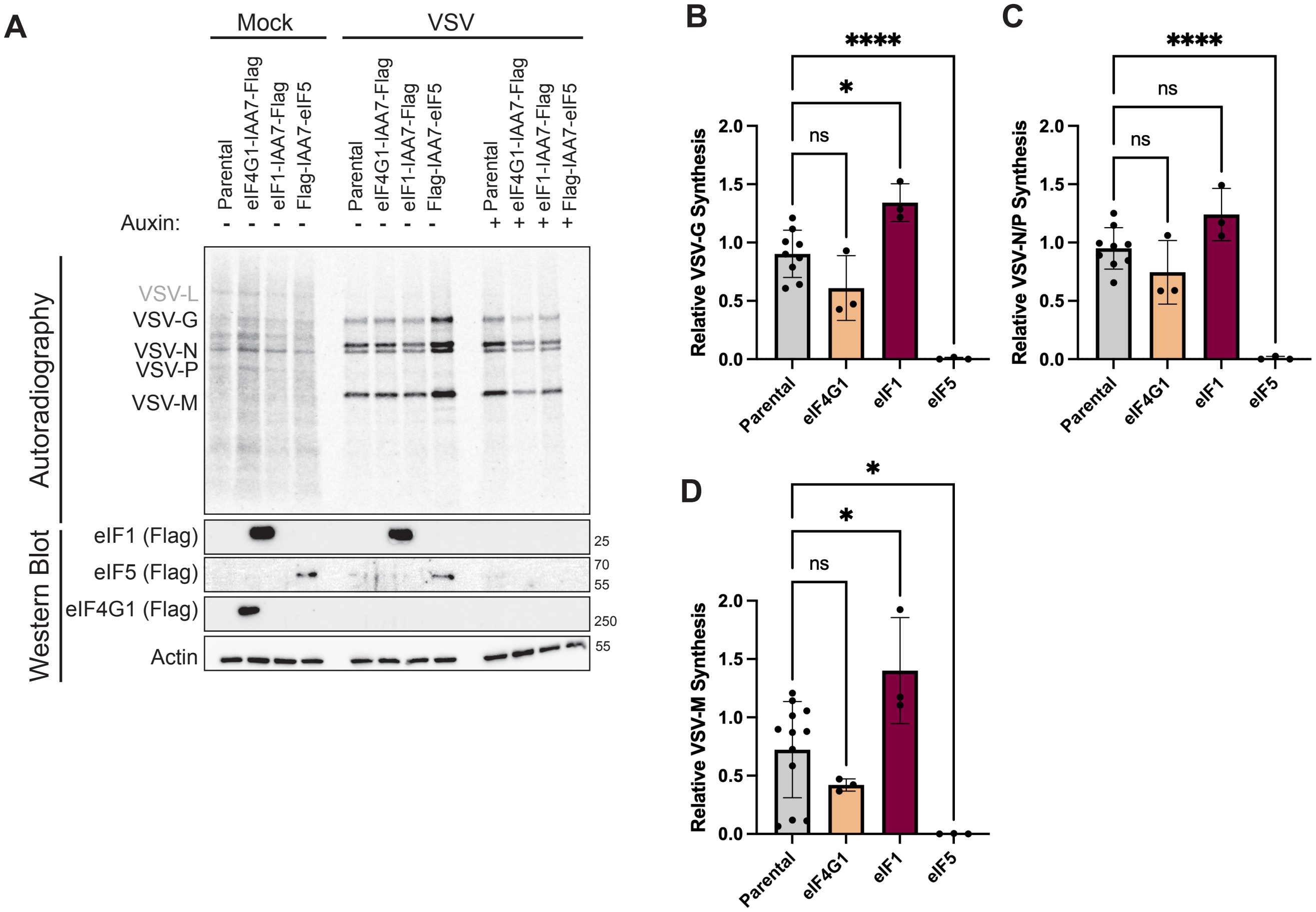

### Supplementary Figure 6

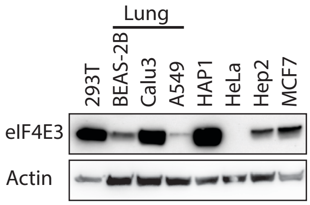
